# Reproducibility test of radiomics using network analysis and Wasserstein K-means algorithm

**DOI:** 10.1101/773168

**Authors:** Jung Hun Oh, Aditya P. Apte, Evangelia Katsoulakis, Nadeem Riaz, Vaios Hatzoglou, Yao Yu, Jonathan E. Leeman, Usman Mahmood, Maryam Pouryahya, Aditi Iyer, Amita Shukla-Dave, Allen R. Tannenbaum, Nancy Y. Lee, Joseph O. Deasy

**Affiliations:** Department of Medical Physics, Memorial Sloan Kettering Cancer Center, New York, NY 10065, USA; Department of Radiation Oncology, Veterans Affairs, James A Haley, Tampa, FL 33612, USA; Department of Radiation Oncology, Memorial Sloan Kettering Cancer Center, New York, NY 10065, USA; Department of Radiology, Memorial Sloan Kettering Cancer Center, New York, NY 10065, USA; Department of Radiation Oncology, Dana Farber Cancer Institute/Brigham and Women’s Hospital, Boston, MA 02189, USA

## Abstract

**Purpose:** To construct robust and validated radiomic predictive models, the development of a reliable method that can identify reproducible radiomic features robust to varying image acquisition methods and other scanner parameters should be preceded with rigorous validation. Due to the property of high correlation present between radiomic features, we hypothesize that reproducible radiomic features across different datasets that are obtained from different image acquisition settings preserve some level of connectivity between features in the form of a network.

**Methods:** We propose a regularized partial correlation network to identify robust and reproducible radiomic features. This approach was tested on two radiomic feature sets generated with two different reconstruction methods from a cohort of 47 lung cancer patients. The commonality of the resulting two networks was assessed. A largest common network component from the two networks was tested on phantom data consisting of 5 cancer samples. We further propose a novel K-means algorithm coupled with the optimal mass transport (OMT) theory to cluster samples. This approach following the regularized partial correlation analysis was tested on computed tomography (CT) scans from 77 head and neck cancer patients that were downloaded from The Cancer Imaging Archive (TCIA) and validated on CT scans from 83 head and neck cancer patients treated at our institution.

**Results:** Common radiomic features were found in relatively large network components between the resulting two partial correlation networks from a cohort of 47 lung cancer patients. The similarity of network components in terms of the common number of radiomic features was statistically significant. For phantom data, the Wasserstein distance on a largest common network component from the lung cancer data was much smaller than the Wasserstein distance on the same network using random radiomic features, implying the reliability of those radiomic features present in the network. Further analysis using the proposed Wasserstein K-means algorithm on TCIA head and neck cancer data showed that the resulting clusters separate tumor subsites and this was validated on our institution data.

**Conclusions:** We showed that a network-based analysis enables identifying reproducible radiomic features. This was validated using phantom data and external data via the Wasserstein distance metric and the proposed Wasserstein K-means method.

## INTRODUCTION

Radiomics enables an in-depth measurement of tumor phenotypes by quantifying imaging signals from radiologic images that can reflect key information of signatures associated with patient outcomes [1, 2]. Recently, the connection of radiomics with machine learning has accelerated the development of new imaging features and radiomic outcomes modeling, showing the potential of using radiomics to build predictive models of individual cancer outcomes [3, 4]. Despite such great progress in radiomics in recent years, however, the development of computational techniques to identify repeatable and reproducible radiomic features remains challenging and relatively retarded [5]. This has led many radiomic models built using a dataset to be unsuccessful in subsequent external validation on independent data [6]. One of the reasons of these consequences is likely due to the susceptibility of radiomic features to image reconstruction and acquisition parameters [7, 8]. Since radiomic features are computed via multiple tasks including imaging acquisition, segmentation, and feature extraction, the selection of parameters present in each step may affect the stability of features computed [9]. As such, prior to model building, development of radiomic features with high repeatability and high reproducibility as well as development of tools that can identify such features is more likely to be urgently needed in the field of radiomics.

In this study, we propose a graph (network)-based computational method that consists of the partial correlation network and graphical lasso (linear absolute shrinkage and selection operator) to identify reliable and reproducible radiomic features, using computed tomography (CT) images in lung cancer patients and phantom data with two reconstruction methods. The proposed method is justified utilizing the *L*^1^-Wasserstein distance (also known as Earth Mover’s distance: EMD) metric measured on radiomic networks. We further propose a novel method, *Wasserstein K-means algorithm*, to cluster samples utilizing the Wasserstein distance as a cost function in the K-means algorithm. This method is tested using CT images in head and neck squamous cell carcinoma (HNSCC) downloaded from The Cancer Imaging Archive (TCIA) and validated using our institution data.

## METHODS

### CT scans in lung cancer patients

This study was approved by internal review board. In total, 47 lung nodules were segmented on contrast enhanced CT of lung cancer patients that were scanned using a GE MEDICAL SYSTEMS scanner and reconstructed with standard and lung convolution kernels. Lung nodules were segmented on scans belonging to the lung reconstruction and were copied over to scans belonging to the standard reconstruction. For each reconstruction method, a set of 132 radiomic features were extracted using the CERR radiomics toolbox [10]. Extracted features fell into three categories: (1) first order statistics, (2) shape-based features, and (3) higher order textures including gray level cooccurrence matrix (GLCM), gray level run length matrix (GLRLM), gray level size zone matrix (GLSZM), neighborhood gray tone difference matrix (NGTDM), and neighboring gray level dependence matrix (NGLDM).

### Phantom data

A multi-material 3D printer (PolyJet Objet 260 Connex 3, Stratasys, Eden Prairie, Minnesota) with Voxel Print software was used for the deposition of droplets of ultraviolet-curable photopolymer resins in a layer-by-layer manner [11]. In this study, two base photopolymer resins were used including TangoPlus (material A) and VeroWhite (material B), which range in attenuation of approximately 65 Hounsfield unit (HU) to 125 HU at 120 kVp. The resolution of the printer is on the order of 48 × 84 × 30 *μ*m. Because the resolution is finer than that of typical CT scanners used at our institution (0.625 × 0.625 × 0.625 mm) [11], multiple resin droplets were mixed in a single voxel to reproduce the contrast differences of tumor specific patterns observed on patient CT scans. To 3D print the fine gradients of tumor intensity patterns, the Floyd Steinberg dithering algorithm was used.

As a first step to produce phantoms, two separate 3D prints were developed. The first 3D print was modeled after a single patient from the RIDER dataset that consisted of patients with non-small-cell lung carcinoma (NSCLC). It was printed to physically visualize the lung vasculature, extent of the tumor and its structure. The second 3D print was designed to capture the morphologic appearance and different intensity patterns seen on CT scans for the single NSCLC and 4 pancreatic ductal adenocarcinoma (PDAC) tumors. Since the surrounding tissue influences the local resolution and noise properties of tumors, the lesions were immersed in a heterogenous background that was modeled after patients with advanced stage hepatic cirrhosis. The area and number of slices of the final 3D print was dictated by the size of the largest tumor.

Prior to dithering, the HU values of each tumor and the background were separately converted to double precision intensity within a range of 0 to 1. The decimal values between the new intensity range dictated the amount of photopolymer material that would be deposited within any given voxel. Then, each slice was super-sampled to the resolution of the 3D printer using the Whittaker–Shannon (SINC) interpolation.

Lastly, each slice was dithered to yield a set of layered raster images using the Floyd-Steinberg dithering algorithm. A single raster layer encodes the spatial allocation of a resin material. Since the 3D printer is capable of depositing 3 different resin materials, three sets of raster files were generated. Within any raster layer, a value of 1 indicates deposition of material A and a value of 0 indicates that no material is deposited. The first set of bitmaps were inverted to mix two materials within a single voxel such that a value of 0 in the first set of bitmaps (material A) has a value of 1 in the second set of bitmaps (material B). Since two materials with opposing densities can generate the desired pattern differences, the third set of bitmaps consisted of all zeros.

After 3D printing, the phantom was scanned sequentially 30 times with a typical abdominal CT protocol using the following parameters: 120 kVp, 280 mA, pitch of 0.984, reconstructed slice thickness of 5 mm, and a reconstruction interval of 5 mm. Images were reconstructed using the filtered back projection algorithm. Two sets of 132 radiomic features were extracted using the CERR toolbox for the standard and lung reconstruction kernels [10].

### CT scans in head and neck cancer patients

For further radiomic network test, pre-treatment CT scans with IV contrast for HNSCC patients were downloaded from the TCIA (http://www.cancerimagingarchive.net/). The data were previously used in our another study [12]. Below we briefly introduce the preprocessing and radiomic feature extraction steps. For more detailed information, see [12]. Available 188 cases were imported into the Eclipse treatment planning system (Varian Medical System, Palo Alto, CA) for segmentation. Before delineating lesions of interest (ROIs), samples that do not meet the inclusion criteria such as primary tumor size and image quality were excluded. For the evaluable scans that fulfill the inclusion criteria, the primary tumor on CT scans was manually delineated by a radiation oncologist and the delineation was independently confirmed by a neuroradiologist. The presence of CT artifacts was further assessed within the primary tumor. Slices with streak artifact were not delineated and excluded from analysis. If the proportion of slices with streak artifact in each tumor is larger than 50% of the number of slices, the case was excluded from the study [13]. This sample quality test resulted in a set of 77 cases with 28 laryngeal, 11 oropharyngeal, and 38 oral cavity tumors. There were 13 HPV-positive and 64 HPV-negative tumors [14].

Radiomic features were extracted using the CERR toolbox on resampled scans at the resolution of 0.6 × 0.6 × 3.5 mm [10]. Due to the effect of sub-sampling slices, which is caused by dental artifacts, two-dimensional radiomic features were extracted: 104 radiomic features including first order statistics and higher order textures were computed from artifact-free slices.

Further feature stability test and volume independent test were performed. For each scan, 100 independent datasets were generated, each of which consisting of 75% of the artifact-free slices and for each dataset 104 radiomic features were recomputed. Features with median coefficient of variation > 0.1 across all samples were removed. Features with high correlations with tumor volume (Spearman’s correlation coefficient > 0.4) were also removed. These two tests resulted in 67 radiomic features that were used for subsequent analysis.

The radiomic analysis results on the TCIA data were validated using an independent dataset with 83 HNSCC patients treated at our institution. The validation cohort consisted of 1 laryngeal tumor, 31 oropharyngeal and 51 oral cavity tumors, all with pre-treatment CT scans with IV contrast. Among 32 patients with laryngeal and oropharyngeal tumors, 27 had HPV-positive and 5 had HPV-negative tumors. However, HPV status for 51 patients with oral cavity tumors was not available since HPV status on oral cavity tumors is not routinely obtained due to the low or rare prevalence of HPV positivity. We utilized the identical radiomic analysis pipeline (including segmentation, preprocessing, feature extraction, and network analysis) used in the analysis of the TCIA data.

### Regularized partial correlation network

For a network representation of radiomic features, we adopted a Gaussian graphical model of partial correlation coefficients in which each connection (link) in a network is represented as a partial correlation coefficient between two radiomic features (nodes) after conditioning on all other available features. This network may be intractably complex including many spurious connections, particularly for data with numerous features. To remove potential false positives and make the network representation more interpretable for a meaningful understanding of the data, the lasso-type regularization (graphical lasso) can be employed in which a tuning parameter *λ* controls the sparsity of the network by shrinking partial correlation coefficients. More specifically, higher *λ* values make the network sparser whereas with lower values a smaller number of connections are removed, making the network denser with likely false-positive connections. To optimize the *λ* value that controls a tradeoff between keeping spurious connections and removing true connections in a network, we use a method that optimizes the fit of the network to the data by minimizing the extended Bayesian information criterion (EBIC).

### Wasserstein distance

The optimal mass transport (OMT) is an active research area with an ever-increasing growth in its application in numerous fields including medical imaging analysis, statistical physics, machine learning, and genomics [15-17]. Here we briefly describe the basic concept of OMT. Let *X* and *Y* denote two probability spaces with measures *μ* and *ν* ∈ *P*(Ω), and let *c*(*x, y*) denote the transportation cost per unit mass from point *x* ∈ *X* to *y* ∈ *Y*. The OMT problem seeks to find a transport map *T*: *X* → *Y* such that the total transportation cost, ∫_*X*_ *c*(*x, T*(*x*))*μ*(*dx*), is minimized. Let us define *P*(Ω) = {*ρ*(*x*): ∫_Ω_ *ρ*(*x*)*dx* = 1, *ρ*(*x*) ≥ 0} where Ω ⊆ ℝ^*m*^. According to the Kantorovich’s relaxed formulation [18], the *L*^1^-Wasserstein distance (EMD) is defined as follows:

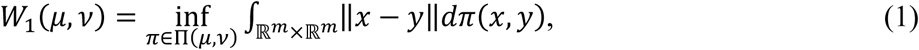

where Π(*μ, ν*) denotes the set of all joint probability measures *π* on Ω × Ω whose marginals are *μ* and *ν*. A computationally more efficient formulation of the Wasserstein distance can be defined in which the flux vector *m*: Ω → ℝ^*m*^ is optimized in the following manner:

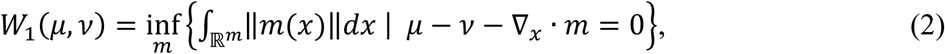

where ‖·‖ is the Euclidean norm.

An alternative graph-theoretic formulation of Eq. (2) to compute the Wasserstein distance on a network is as follows:

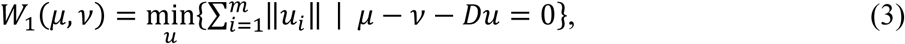

where *m* is the number of edges in a network, *u*_*i*_ are fluxes on the edges, and *D* is an incidence matrix with rows and columns indexed by the nodes and edges in the network such that every entry (*i, k*) is set to 1 if the node *i* is assigned to be the head of the edge *k* and is set to -1 if it is the tail of *k*. Using Eq. (3), we can compute the Wasserstein distance between two samples on a connected component of partial correlation network consisting of radiomic features. We used the CVX toolbox to optimize the OMT problem.

### Wasserstein K-means algorithm

The K-means algorithm is one of the most commonly used clustering algorithms that partition a given set of samples into *c* clusters, by minimizing the within-cluster sum of squares:

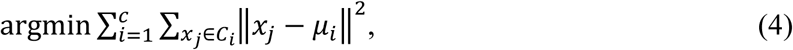

where *μ*_*i*_ is the mean of samples in cluster *C*_*i*_. Here we propose a novel clustering method in which the cost function in the conventional K-means algorithm is replaced with the Wasserstein distance metric as shown in the following equation:

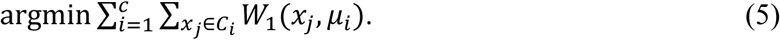

In this approach, during the process of K-means clustering, distances from each sequentially updated centroid in *c* clusters to samples on a fixed optimized radiomic network are computed and clustering of the samples is performed such that the within-cluster sum of Wasserstein distance is minimized. The K-means algorithm coupled with the Wasserstein distance intuitively enables us to cluster samples in a network form of data. We call this method a *Wasserstein K-means algorithm*.

## RESULTS

### Regularized partial correlation network

Two different reconstruction kernels (lung and standard) were applied to CT scans for a cohort with 47 lung nodules, generating a set of 132 radiomic features for each reconstruction method. For each set of features, a radiomic network was constructed using regularized partial correlation coefficients coupled with EBIC. As a result, two different radiomic networks were constructed. To remove potential false-positive relationships between nodes (features) in the networks, connections with partial correlation coefficients < 0.2 were further removed, resulting in more network components (“islands”). Hereafter, we define a component as a connected set of nodes in a network.

Figure 1 shows the two largest network components for each reconstruction method with the number of links ≥ 9: Figures 1A and 1B resulted from the standard reconstruction method whereas Figures 1C and 1D resulted from the lung reconstruction method. In comparison of Figures 1A and 1C, 10 radiomic features were common, including First order statistics: entropy (10), Shape: surface to volume ratio (39), GLRLM: run entropy (74), NGTDM: coarseness (82), contrast (83), strength (86), NGLDM: dependence count non-uniformity normalized (98), dependence count entropy (101), dependence count energy (102), and GLSZM: size zone entropy (118). In comparison of Figures 1B and 1D, 5 radiomic features were common including GLRLM: high gray level run emphasis (69), short run high gray level emphasis (80), NGLDM: high grey level count emphasis (90), GLSZM: high gray level zone emphasis (111), and small area high gray level emphasis (113). In the similarity test of network components using the hypergeometric distribution, statistically significant p-values were obtained with 2.3×10^−8^ between Figures 1A and 1C and 4.2×10^−4^ between Figures 1B and 1D, showing the strong reproducibility of these radiomic features between different reconstruction methods. Note that the similarity of network topologies was not considered.

**Figure 1.**
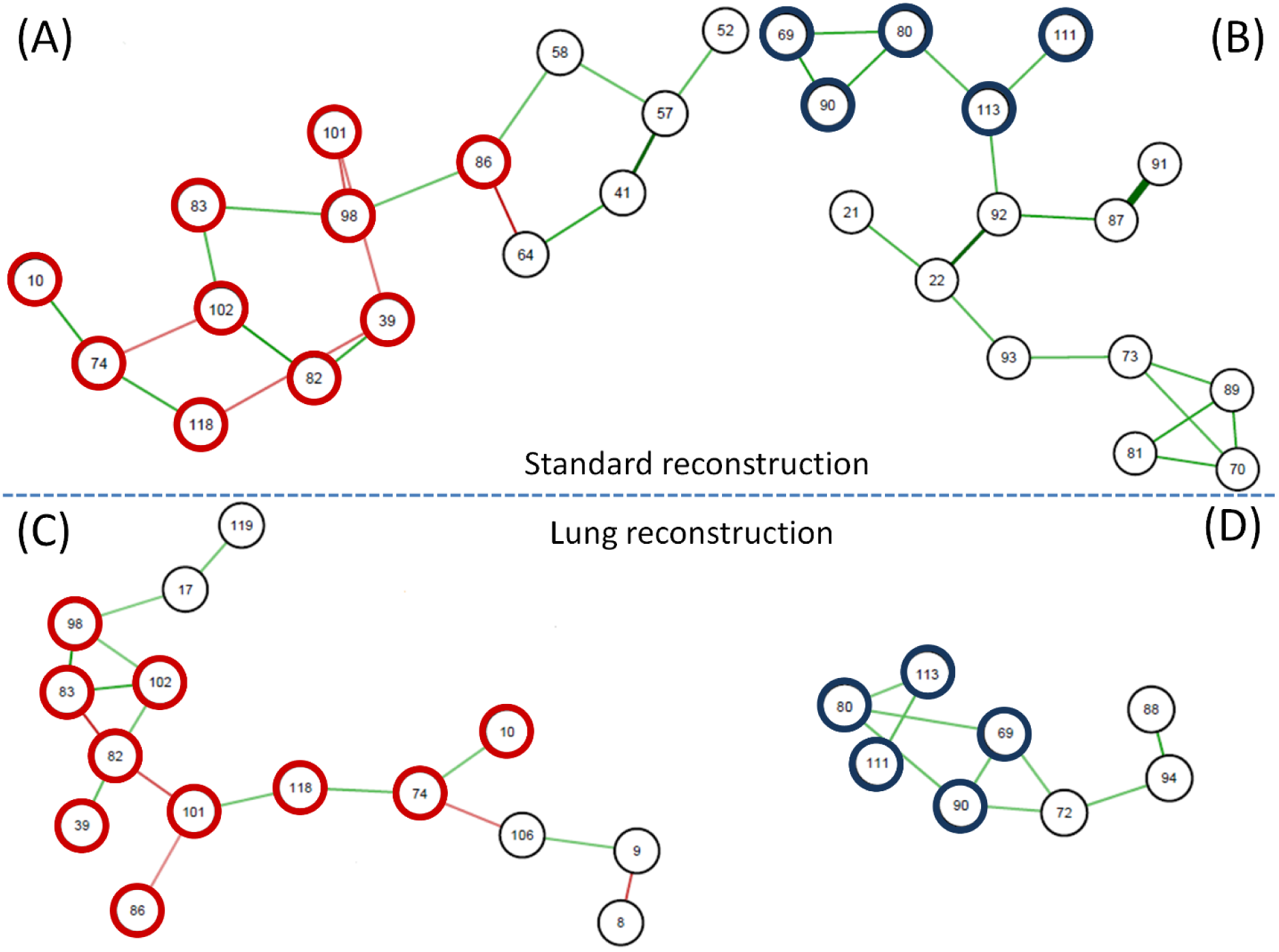
The two largest network components from the standard reconstruction kernel (A and B) and from the lung reconstruction kernel (C and D). The thick circles indicate the common radiomic features between two network components (each one from two different reconstruction kernels). The numbers in the circles indicate the order of 132 features in our data.

### Validation on phantom data

To further test the reproducibility of these features on phantom data, the two largest components (each one from two different reconstruction kernels) shown in Figures 1A and 1C were merged, making a larger network with 20 unique radiomic features. As previously described, 5 phantoms were generated in this study and 10 sets of radiomic features (2 sets for each phantom) were extracted using the same two reconstruction kernels (standard and lung). Using Eq. (3), the Wasserstein distance was computed on the merged network between the two sets of radiomic features (standard and lung) of each sample (Figure 2). The average Wasserstein distance on the 5 phantoms was 0.21. It is likely that if those 20 radiomic features in the merged network are reliable and stable, the Wasserstein distance using random radiomic features on the same network will be larger than 0.21. With this hypothesis, we randomly selected 20 features from the available 132 radiomic features (randomly assigning the 20 features to the 20 nodes in the merged network) and computed an average Wasserstein distance of 5 phantoms. The whole process was repeated 1000 times. An average Wasserstein distance of 5 phantoms after the 1000 iterations was 0.32 and for only 49 times out of 1000 the average Wasserstein distance on the random simulation test was less than 0.21, implying the stability of the 20 radiomic features in the merged network (Figure 3).

**Figure 2.**
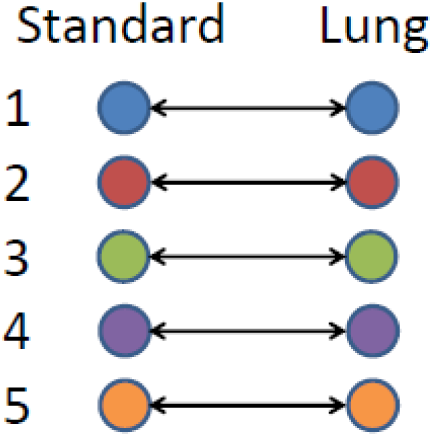
For each phantom, the Wasserstein distance was computed between a set of features of standard reconstruction and a set of features of lung reconstruction on a network that was constructed using lung cancer data.

**Figure 3.**
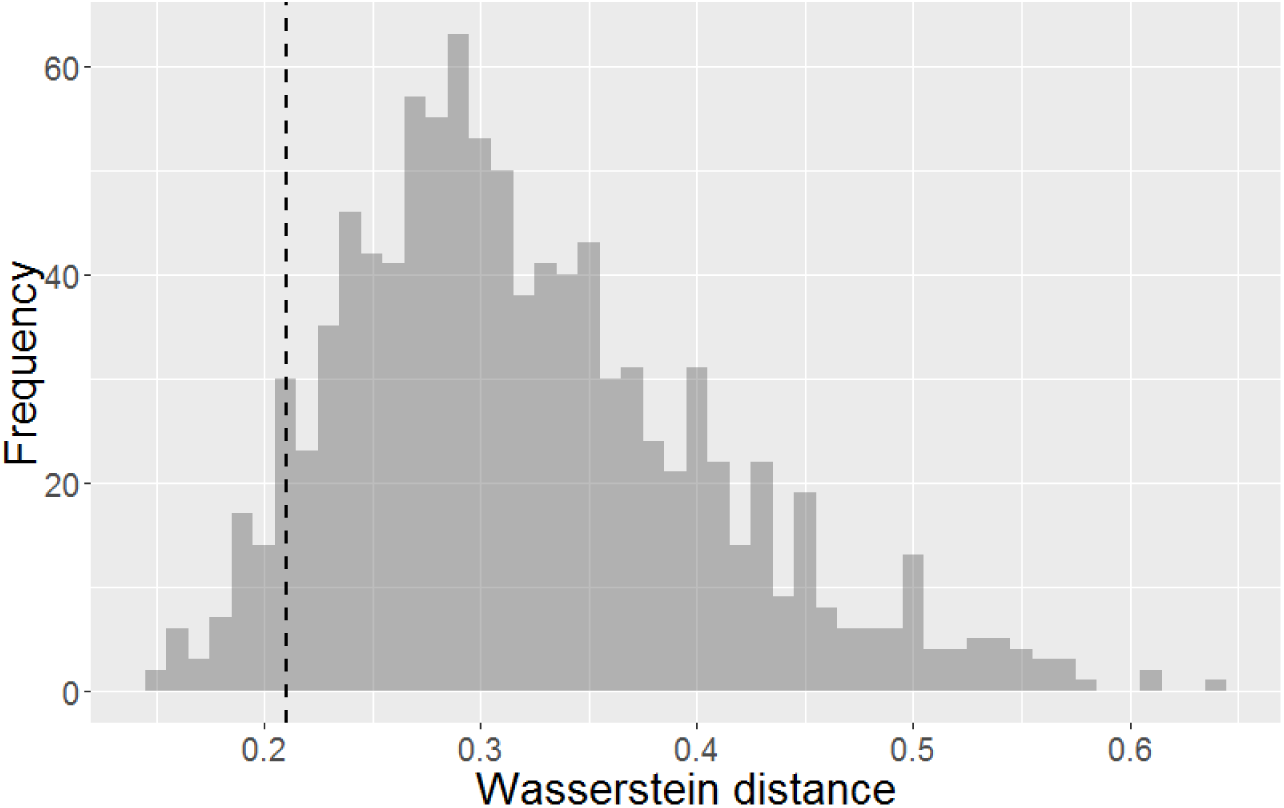
For each phantom, the Wasserstein distance was computed between a set of features of standard reconstruction and a set of features of lung reconstruction on the network that was constructed using lung cancer data, by randomly selecting 20 radiomic features out of 132 and randomly assigning them to the nodes in the network. After each iteration, an average Wasserstein distance for the 5 phantoms was computed. This histogram shows the 1000 average values after the 1000 simulation tests. The dotted vertical line indicates 0.21 that is an average Wasserstein distance of the 5 phantoms computed on the original network that was constructed using lung cancer data.

### Wasserstein K-means clustering

Using TCIA head and neck cancer data that consisted of 77 samples and 67 radiomic features, a regularized correlation network was constructed. After applying the EBIC and cutting the links with a partial correlation coefficient < 0.2, network components were generated. The three largest components with the number of links ≥ 9 were chosen for further analysis, which consisted of 26 radiomic features (Figure 4). Using the Wasserstein K-means algorithm, clustering was performed. Based on our previous study [12], K=2 was chosen. During the K-means clustering process, the Wasserstein distance was computed for each component and the three Wasserstein distances were averaged. For visualization purpose, principal component analysis (PCA) was carried out on the 26 radiomic features. Figure 5 shows the clustering results visualized using the first two principal components.

**Figure 4.**
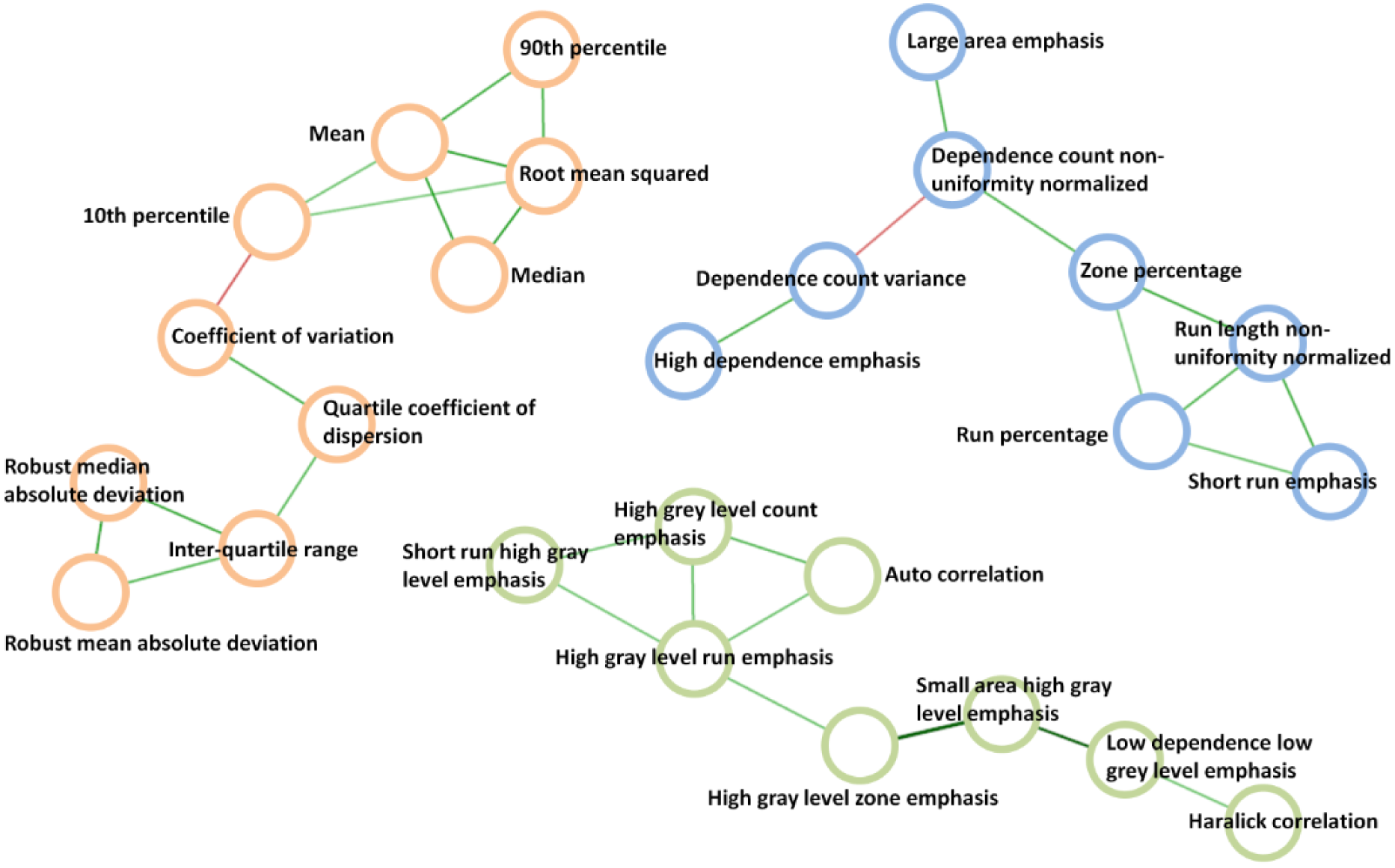
The three largest components of partial correlation network that resulted from the TCIA head and neck cancer data.

**Figure 5.**
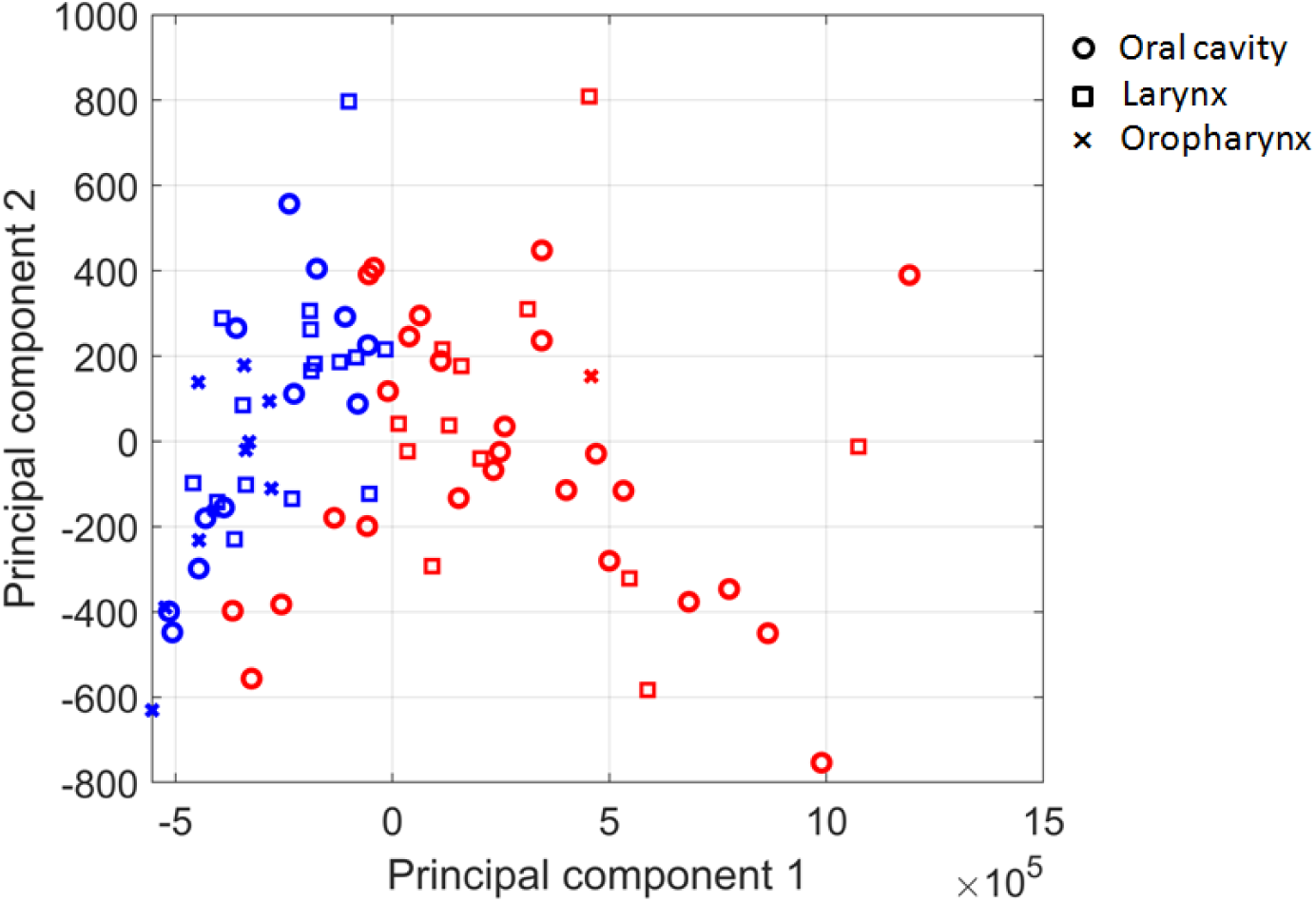
The Wasserstein K-means clustering algorithm was performed on TCIA head and neck cancer data. For visualization purpose, principal component analysis was performed and the final clustering results were projected to the first two principal components. The blue and red colors indicate the two different clusters.

A significant difference in tumor subsite was found between the two radiomic clusters with extended Fisher’s exact test p=0.0063. One cluster (red) had 1 oropharyngeal, 26 oral cavity, and 12 laryngeal tumors whereas the other cluster (blue) had 10 oropharyngeal, 12 oral cavity, and 16 laryngeal tumors and this cluster was significantly enriched for HPV-positive tumors with extended Fisher’s exact test p=0.0012. These results were similar to what we previously reported in another study [12].

For validation, the same 26 radiomic features were extracted from CT scans for 83 HNSCC patients treated at our institution and PCA was performed on those 26 features. Figure 6 shows the mapping results using the first two principal components. Similarly, oropharyngeal tumors were clustered and separated from oral cavity tumors. Using a classifier with the dotted line boundary, the accuracy of classifying oropharyngeal tumors was 93.6% (29/31) and the accuracy of classifying oral cavity tumors was 72.6% (37/51).

**Figure 6.**
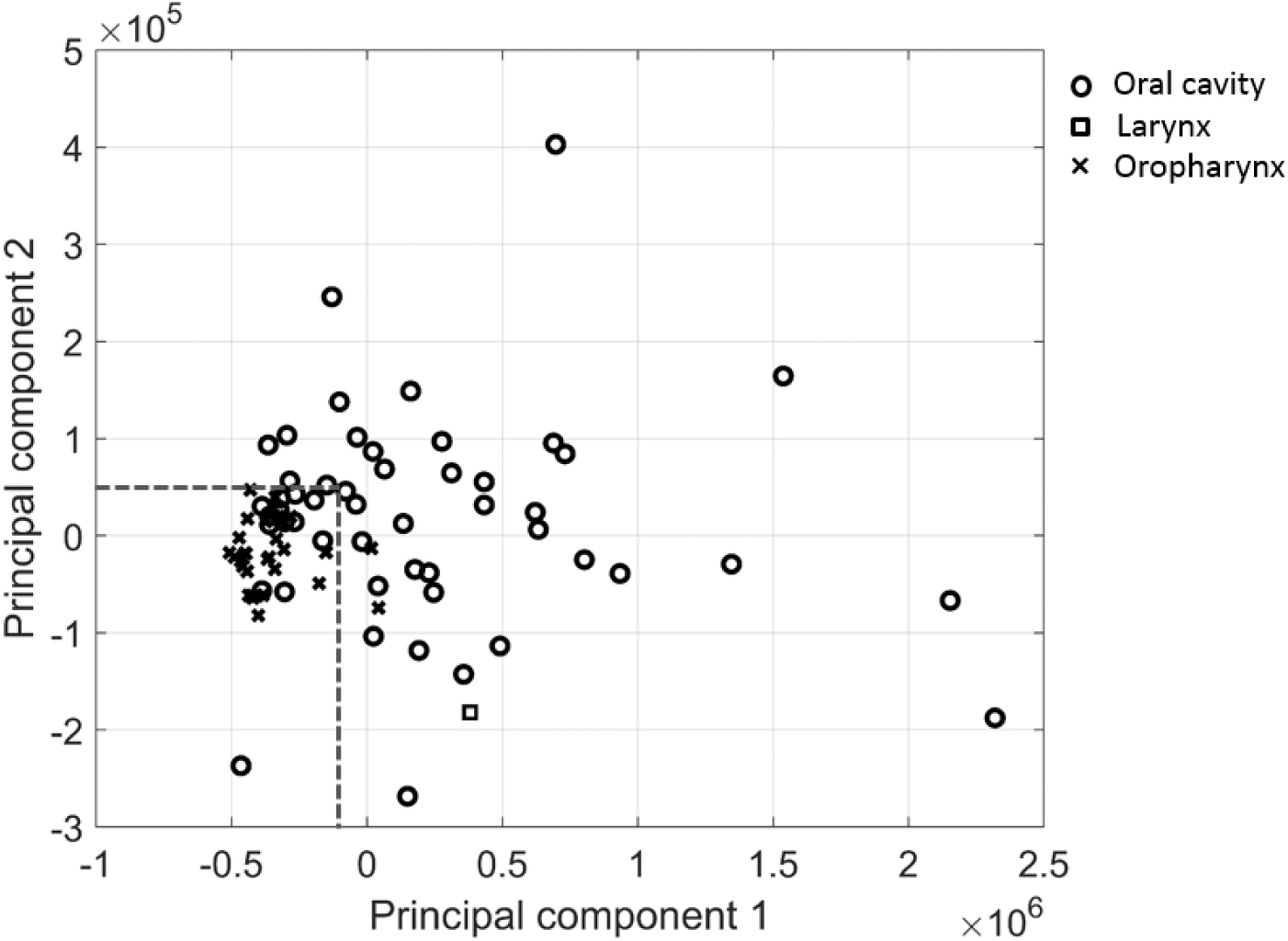
For validation, the same 26 radiomic features were extracted from CT scans for 83 head and neck cancer patients treated at our institution. Principal component analysis was then carried out on the data. This scatter plot shows the mapping results using the first two principal components.

## CONCLUSION

In this study, we showed the potential of using a network-based approach to identify reliable and reproducible radiomic features that are robust to different image reconstruction methods. This was tested on cancer and phantom data using the Wasserstein distance metric designed to compute the dissimilarity of samples on a network. We further proposed a Wasserstein K-means algorithm to cluster samples by replacing the cost function of Euclidean distance in the conventional K-means algorithm with the Wasserstein distance metric. The clustering results on TCIA data were validated using independent data. The OMT is an active research topic. Applying the OMT coupled with the network analysis to radiomics could provide a powerful tool to identify reproducible radiomic features as well as develop robust prediction models.

## Acknowledgements

This research was funded in part through National Institutes of Health/National Cancer Institute Cancer Center Support grant P30 CA008748.

## References

1. Gillies RJ, Kinahan PE, Hricak H: Radiomics: Images Are More than Pictures, They Are Data. Radiology 2016, 278:563–577.

2. Aerts HJ, Velazquez ER, Leijenaar RT, Parmar C, Grossmann P, Carvalho S, Bussink J, Monshouwer R, Haibe-Kains B, Rietveld D et al: Decoding tumour phenotype by noninvasive imaging using a quantitative radiomics approach. Nat Commun 2014, 5:4006.

3. Giraud P, Giraud P, Gasnier A, El Ayachy R, Kreps S, Foy J-P, Durdux C, Huguet F, Burgun A, Bibault J-E: Radiomics and Machine Learning for Radiotherapy in Head and Neck Cancers. Front Oncol 2019, 9:174.

4. Folkert MR, Setton J, Apte AP, Grkovski M, Young RJ, Schöder H, Thorstad WL, Lee NY, Deasy JO, Oh JH: Predictive modeling of outcomes following definitive chemoradiotherapy for oropharyngeal cancer based on FDGPET image characteristics. Phys Med Biol 2017, 62(13):5327–5343.

5. Traverso A, Wee L, Dekker A, Gillies R: Repeatability and Reproducibility of Radiomic Features: A Systematic Review. Int J Radiat Oncol Biol Phys 2018, 102(4):1143–1158.

6. Virginia BM, Laura F, Silvia R, Roberto F, Francesco F, Eva H, Charles F, Samy A, Stefan M, Jean-Charles S et al: Prognostic value of histogram analysis in advanced non-small cell lung cancer: a radiomic study. Oncotarget 2018.

7. Zhao B, Tan Y, Tsai W-Y, Qi J, Xie C, Lu L, Schwartz LH: Reproducibility of radiomics for deciphering tumor phenotype with imaging. Sci Rep 2016:srep23428.

8. Rizzo S, Botta F, Raimondi S, Origgi D, Fanciullo C, Morganti AG, Bellomi M: Radiomics: the facts and the challenges of image analysis. Eur Radiol Exp 2018, 2:36.

9. Park JE, Park SY, Kim HJ, Kim HS: Reproducibility and Generalizability in Radiomics Modeling: Possible Strategies in Radiologic and Statistical Perspectives. Korean J Radiol 2019, 20(7):1124–1137.

10. Apte AP, Iyer A, Crispin-Ortuzar M, Pandya R, van Dijk LV, Spezi E, Thor M, Um H, Veeraraghavan H, Oh JH et al: Technical Note: Extension of CERR for computational radiomics: A comprehensive MATLAB platform for reproducible radiomics research. Med Phys 2018, 45(8):3713–3720.

11. Obuchowski NA, Barnhart HX, Buckler AJ, Pennello G, Wang XF, Kalpathy-Cramer J, Kim HJ, Reeves AP: Statistical issues in the comparison of quantitative imaging biomarker algorithms using pulmonary nodule volume as an example. Stat Methods Med Res 2015, 24(1):107–140.

12. Katsoulakis E, Yu Y, Apte AP, Leeman JE, Katabi N, Morris L, Deasy JO, Chan TA, Lee NY, Riaz N et al: Radiomic analysis of head and neck squamous cell carcinoma (HNSCC) identifies subtypes that correlate with distinct molecular and microenvironmental characteristics of tumors. Under review 2019.

13. Ger RB, Craft DF, Mackin DS, Zhou S, Layman RR, Jones AK, Elhalawani H, Fuller CD, Howell RM, Li H et al: Practical guidelines for handling head and neck computed tomography artifacts for quantitative image analysis. Comput Med Imaging Graph 2018, 69:134–139.

14. Nulton TJ, Olex AL, Dozmorov M, Morgan IM, Windle B: Analysis of The Cancer Genome Atlas sequencing data reveals novel properties of the human papillomavirus 16 genome in head and neck squamous cell carcinoma. Oncotarget 2017, 8(11):17684–17699.

15. Villani C: Optimal transport: old and new: Springer Science & Business Media; 2008.

16. Chen Y, Cruz FD, Sandhu R, Kung AL, Mundi P, Deasy JO, Tannenbaum A: Pediatric sarcoma data forms a unique cluster measured via the earth mover’s distance. Scientific reports 2017, 7:7035.

17. Pouryahya M, Oh JH, Javanmard P, Mathews J, Belkhatir Z, Deasy J, Tannenbaum AR: A Novel Integrative Multiomics Method Reveals a Hypoxia-Related Subgroup of Breast Cancer with Significantly Decreased Survival. bioRxiv 2019.

18. Kantorovich L: On the transfer of masses. Dokl Akad Nauk SSSR 1942, 37:227–229.

